# Multi-Scale Tri-Modal Histology Dataset Integrating Tumor Morphology, Immune Patterns, and Clinical Outcomes

**DOI:** 10.64898/2026.05.15.725535

**Authors:** Kyeong Joo Jung, Jianwei Qiu, Sanghee Cho, Elizabeth McDonough, Chrystal Chadwick, Soumya Ghose, Robert West, James D. Brooks, Fiona Ginty, Raghu Machiraju, Parag Mallick

## Abstract

Accurate prognostic assessment of prostate cancer (PCa) requires an integrated understanding of tissue morphology-encompassing cell structure, glandular architecture, and tissue organization-and the immune environment. We present **Prostate-TriMod**, a novel tri-modal histology dataset designed to integrate high-resolution visual morphology with spatial tissue maps, immune infiltration patterns, and clinical outcomes. This dataset, generated from the Cell DIVE™ multiplexed imaging platform, consists of three synchronized modalities: (1) multiscale virtual H&E tiles (224px, 256px, 512px, and 2040px) providing visual morphological context, (2) spatial tissue maps identifying cancerous/non-cancerous epithelial cells, stroma and immune cell populations (via TOPAZ and CAT models), and (3) text captions generated from single-cell data and patterns. The dataset includes comprehensive clinical annotations, including Grade Groups and biochemical recurrence (BCR) status. By providing high-fidelity alignment between visual features, spatial tissue maps, and textual descriptions, Prostate-TriMod empowers the development of advanced multimodal AI frameworks. We expect this resource to support reuse in multimodal representation learning, spatial analysis, and benchmarking studies that link histology morphology and immune context to clinical outcomes in prostate cancer.

## Background & Summary

Prostate cancer (PCa) is the second most commonly diagnosed cancer and the fifth leading cause of cancer death in men worldwide^1,2^. There is a large discrepancy between the number of PCa cases diagnosed and the number of men who die of their disease. For example, in the US in 2026, an estimated 333,830 new PCa cases will be diagnosed, while only 36,320 will die of their disease^3^. It is essential to achieve accurate prognostic assessment to build personalized therapeutic strategies.

Prostate cancer diagnosis and prognosis primarily rely on the manual Gleason scoring system, a histology-based metric that evaluates glandular morphological patterns^4^. This system assigns grades from 1 to 5 based on the morphological patterns of the tumor. Since prostate tumors often exhibit heterogeneous morphology, the final Gleason Score (GS) is determined by summing the two most prevalent patterns^5^. These scores (ranging from 2 to 10) are further categorized into five Grade Groups^6^. While Grade Group 1 (GS ≤ 6) typically represents indolent disease with favorable outcomes, Grade Groups 2 through 5 (GS 7 to 10) indicate a progressive increase in the risk of metastatic spread, recurrence after local therapy, and PCa-specific mortality.

In recent years, there have been milestones in predicting Gleason Grade Groups and biochemical recurrence using deep learning-based artificial intelligence (AI) models, particularly those utilizing histopathological images^7,8^. However, the development of multimodal foundation model is currently constrained by limitations in dataset composition and annotation granularity in existing public datasets. Training and validating such multimodal models require datasets that jointly represent morphology, cell-type context, and interpretable textual descriptors at matched spatial scales.

Commonly used publicly available datasets, such as TCGA-PRAD^9^ and PANDA^7^, rely on Hematoxylin and Eosin (H&E) stained images paired with slide-level labels, and do not capture the cellular interactions between the tumor and tumor microenvironment (TME) (Table 1). Slide-level labels of Gleason Grade Groups fail to capture cellular, immunologic, and stromal features and context in the histological images, leaving these features to be imputed from genomic deconvolution bulk transcriptomic data rather than from direct spatial observation. Diagnostic datasets like PANDA focus heavily on Gleason grading but provide no information regarding the immune microenvironment. While the PESO dataset^10^ offers tile-level annotations for cancerous and non-cancerous epithelium, it lacks comprehensive cell-type maps (including stroma and immune populations) and clinical outcome data which is needed for prognostic modeling. Similarly, the HelsinkiProstate dataset^11^ incorporates cell-type maps but does not provide descriptive text captions to assist in multimodal reasoning. Even broader efforts like Quilt-1M^12^ and ProvPath^13^ utilize AI-generated reports or pathology reports that lack structured, unified format and precise cell-type maps necessary for modeling complex tumor-immune interactions.

**Table 1.** Comparison of the Prostate-TriMod dataset with public benchmarks. (Tri-modal: Integration of Virtual H&E, Cell-type Maps, and Text Captions. C / T / S: Hierarchical annotations provided at the Cell (coordinates and phenotypes), Tile (local spatial contexts at 224px, 256px, and 512px), and Slide levels (whole-tissue master records at 2040px).)

| Dataset | Modality | Annotation Level | Cell type Map | Text Caption | Immune Context | Clinical outcomes |
| --- | --- | --- | --- | --- | --- | --- |
| TCGA-PRAD (Prostate) | H&E + Omics | Slide-level | No | No | Via deconvolution | Various |
| PANDA (Prostate) | H&E | Slide/Tile-level | No | No | No | Grade |
| PESO (Prostate) | H&E | Tile-level | No | No | No | No |
| Quilt-1M (Multi-organ) | H&E + Text | Slide-level | No | Generic Caption from literature<br>AI-corrected from Youtube | Limited | No |
| ProvPath (Multi-organ) | H&E + Text | Slide-level | Yes | WSI report pairs using AI | No | Various |
| HelsinkiProstate (Prostate) | H&E | Tile-level | Yes | No | No | No |
| Prostate-TriMod (Prostate) | Tri-modal | C/T/S | Yes | Captions based on single cell data | Yes | Recurrence, Grade |

To bridge these gaps, we introduce **Prostate-TriMod**, a novel Tri-modal Prostate Cancer Histology Dataset—integrating high-resolution H&E images (20x), pixel-level cell-type maps, and multiscale structured textual captions—with the generation workflow illustrated in Figure 1. The dataset presents four features designed to enhance multimodal AI training and biological discovery.

**Figure 1.**
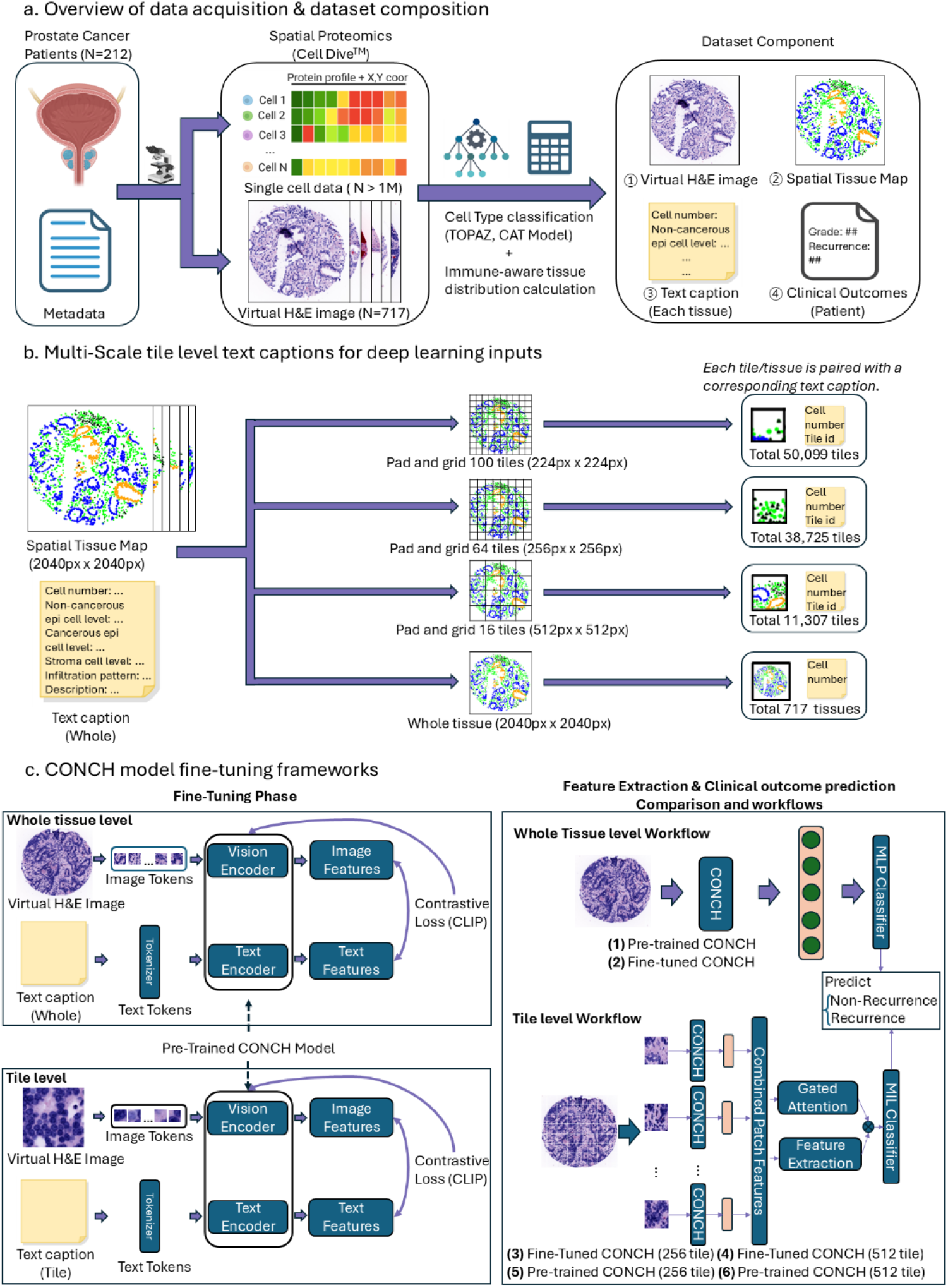
Overview of data acquisition, multi-scale dataset composition, and model fine-tuning frameworks for prostate cancer biochemical recurrence prediction. (a) Spatial single cell proteomics data (over 1 million cells) and virtual H&E images (717 tissue cores) are acquired from the 212 prostate cancer patients. Using the cell type classification models (TOPAZ and CAT models) and immune aware feature calculations, 4 different types of multi-modal data types are generated: (1) Virtual H&E image, (2) Tissue map, (3) Comprehensive text caption including immune infiltration context, and (4) clinical outcomes (biochemical recurrence (yes/no), and Grade Group). (b) 4 different scale text captions are generated to manage with diverse sizes of deep learning inputs. Each tile (224px; 100tiles/tissue, 256px; 64 tiles/tissue, 512px tiles/tissue), and whole tissue(2040px) are paired with corresponding text captions. Prior to tiling, padding was done on the tissues. (c) CONCH model fine-tuning framework to present the validation of the data quality. The fine-tuning process utilizes contrastive learning (CLIP) to align visual features with the text features. Performance of clinical outcome prediction was done to show that the performance includes after the fine-tuning under 4 different conditions: (1) pre-trained CONCH to tissue image, (2) fine-tuned CONCH to tissue image using multi-layer perceptron classifier, (3-4) fine-tuned tile-level CONCH (256px and 512px resolution) using multiple instance learning (MIL) classifier, (5-6) pre-trained tile-level CONCH (256px and 512px resolution) using multiple instance learning (MIL) classifier.

- We provide **multiscale structured text captions** generated across multiple tiles and whole tissue resolutions (224px, 256px, 512px, and 2040px). Unlike existing datasets that rely on noisy natural language, this multiscale approach allows models to capture pathological features ranging from localized cellular distributions to broad architectural patterns within a fixed 20x magnification.
- Our dataset features **high-resolution spatial tissue maps** that are precisely aligned with virtual H&E images. By mapping epithelial, stromal, and immune cell distributions, we provide a structured biological context that enables researchers to investigate the spatial organization of the tumor microenvironment without loss of morphological context.
- Our dataset is uniquely **immune-aware, integrating spatial immune infiltration patterns** directly into the text captions. This provides direct observation of the immune microenvironment, a factor in PCa progression^14,15^ that is often overlooked in existing resources (Table 1).
- We ensure **high clinical relevance** through outcome integration, specifically linking our dataset to BCR and Grade Groups. By bridging morphological features with immune characteristics and patient outcomes, this dataset provides a robust benchmark for developing predictive models that can identify high-risk patients. Predictive modeling experiments are included solely to validate the consistency and utility of the dataset, not to propose new modeling approaches.

Through the technical validation presented in this study, including multiscale consistency (Figure 4) and correlation with clinical progression (Figure 5 and 6), we demonstrate that this dataset provides a multimodal dataset of prostate pathology.

Collectively, **Prostate-TriMod** complements existing resources prostate resources by unifying morphology, immune context, text captions, and clinical outcomes in a single, well-documented corpus. By synchronizing these modalities across local tiles and whole-tissue contexts, the dataset preserves biologically representative tumor–immune architecture while remaining directly usable for modern multimodal learning paradigms, such as vision-language models (VLMs) and multiple instance learning (MIL) pipelines. We anticipate that Prostate-TriMod will serve as a rigorously curated foundation for the community to explore immune-aware multimodal learning and risk-stratification research in prostate cancer, and to support the development and benchmarking of clinically relevant computational workflows.

## Methods

### Overview of the Data Generation Pipeline

The generation of the Prostate-TriMod followed a systematic pipeline designed to integrate morphological, cellular, and textual information. As illustrated in Figure 1, the workflow consists of four primary stages: (1) tissue specimen acquisition and digitization for spatial proteomics via the Cell DIVE™ platform (Leica Microsystems)^16^, (2) cell type classification mapping using TOPAZ^15^ and CAT^17^ models from single-cell data, (3) multiscale image pre-processing and tiling, and (4) rule-based textual caption generation based on quantitative single-cell metrics.

### Patient Cohort and Clinical Characteristics

The study utilized formalin fixed paraffin embedded tissues from a cohort of prostate cancer patients from the Stanford University who underwent radical prostatectomy between 1996 and 2009. All patients signed informed consent that states explicitly that their tissues can be used for genomic and proteomic research and linked to their de-identified clinical data. The protocol was approved with ongoing renewals by the Stanford University Institutional Review board (IRB# 11612). The initial sample consisted of 3 tissue microarrays (TMAs) consisting of 754 cores from 234 patients. After applying rigorous quality control (QC) exclusions and during tissue core removal from TOPAZ workflow, the final analytical dataset was refined to 717 cores from 212 patients. With a median of 9 years of clinical follow-up, 42 patients experienced biochemical recurrence following surgery while the remainder had no evidence of recurrence. The comprehensive clinical and demographic characteristics of the cohort—including age, ethnicity, follow-up time, and Grade Groups—are summarized in Figure 2. These samples provided the foundational material for the subsequent multiplexed imaging and spatial proteomic pipeline illustrated in Figure1.

**Figure 2.**
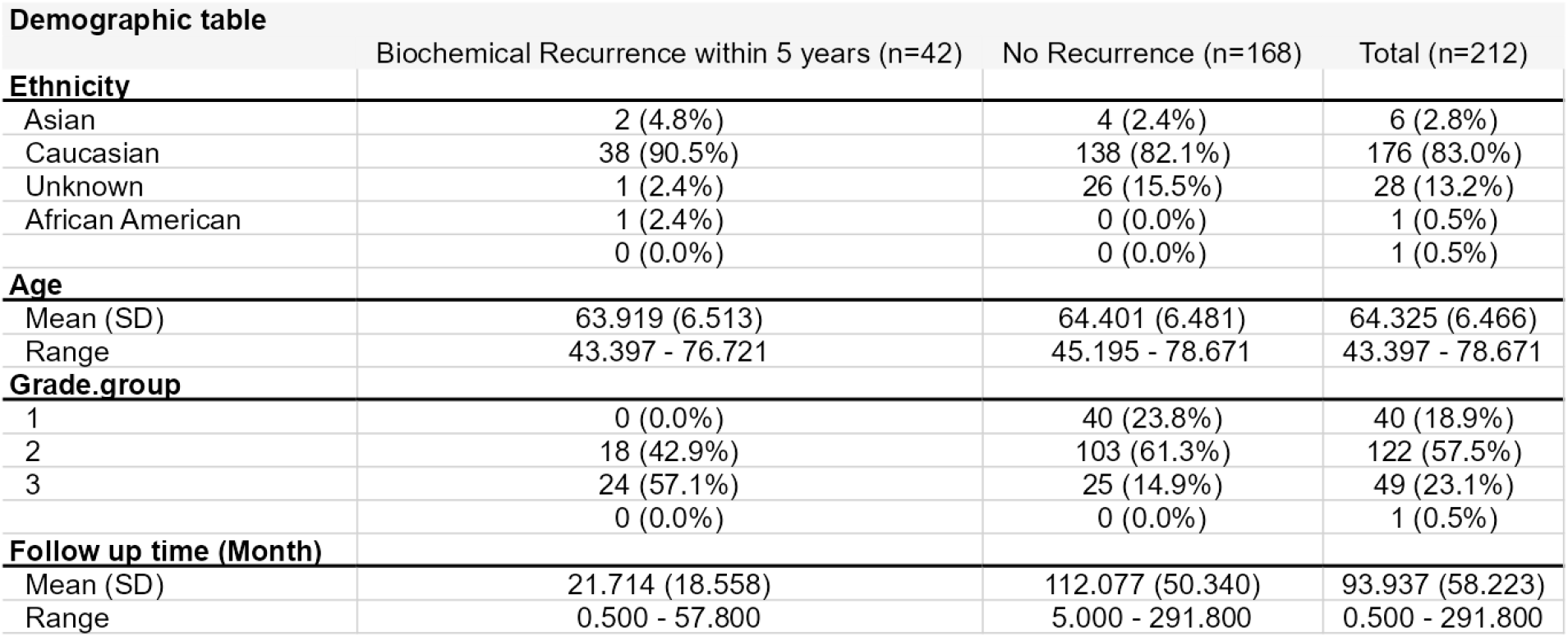
Demographic table of the cohort.

### Multiplexed Imaging and Virtual H&E Generation

High-dimensional spatial proteomic data were acquired using the Cell DIVE™ multiplexed imaging platform (Leica Microsystems). The imaging panel targeted 12 protein markers essential for characterizing the prostate tumor microenvironment (TME):

- Basal and Epithelial Segmentation: CK5, p63, NaKATPase, S6, panCK-PCK26, and panCK-AE1.
- Immune Lineage Identification: CD3, CD4, CD8, CD68, and FOXP3.
- Cancer cell marker and Nuclear stain: AMACR and DAPI.

To facilitate pathological review and maintain alignment with traditional workflows, virtual H&E (vH&E) images (Figure 3a) were synthesized by merging DAPI (nuclei) and autofluorescence (background/stroma) channels at 20x magnification. This mapping process provides a pixel resolution of approximately 0.25 μm/pixel, ensuring that the images are fully registered with the underlying single-cell data.

**Figure 3.**
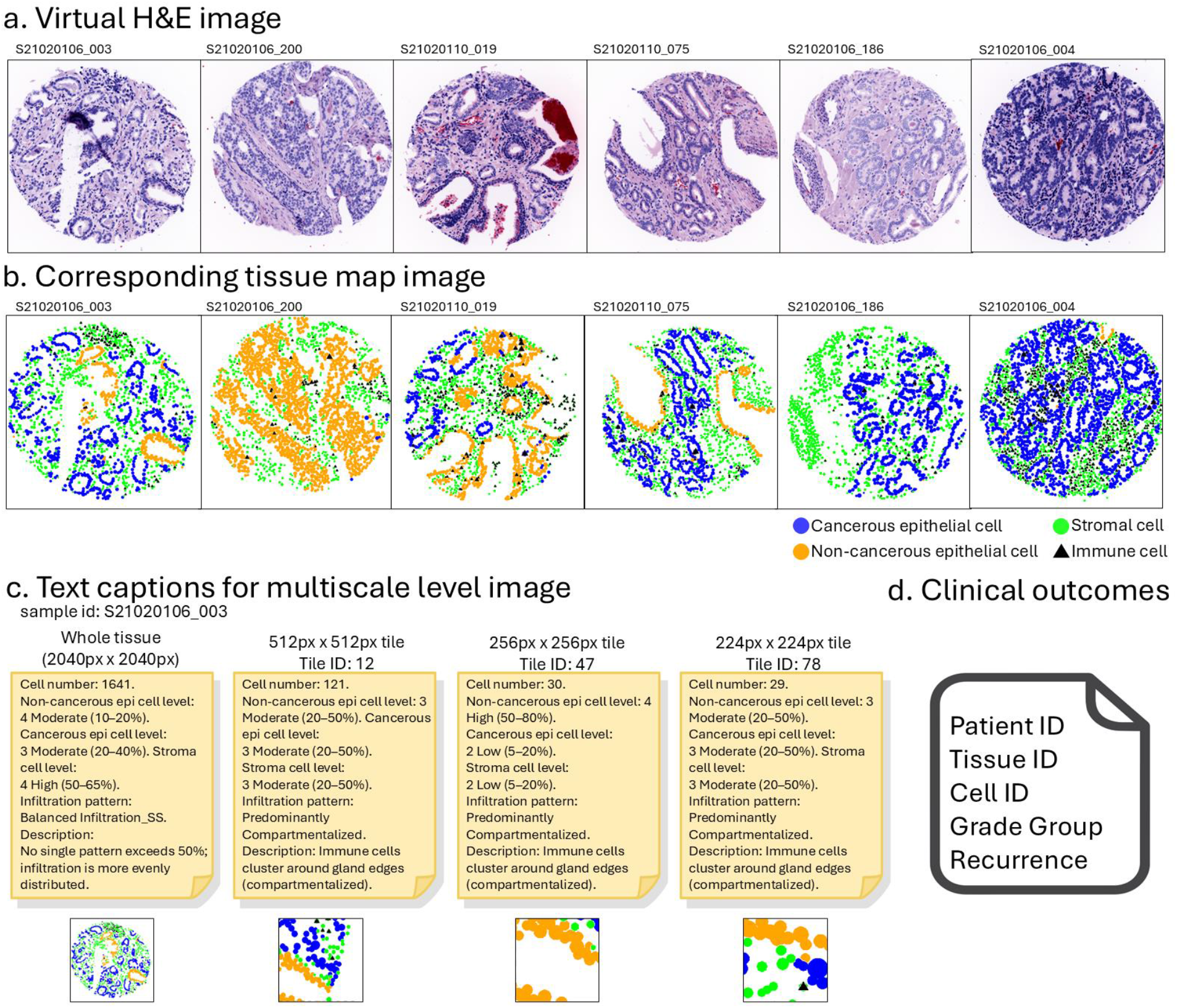
Examples of the Prostate-TriMod. 6 different (a) prostate cancer virtual H&E (vH&E) images at fixed 20x magnification, (b) corresponding tissue maps (including cancerous epithelial cell, non-cancerous epithelial cell, stromal cell, immune cell), (c) text captions under different scales (2040px/512px/256px/224px) which contains number of cells, distribution level, infiltration pattern, and corresponding infiltration pattern description. (d) Clinical outcomes of Grade Group, and biochemical recurrence events along with metadata including patient ID, tissue ID, and cell ID.

**Figure 4.**
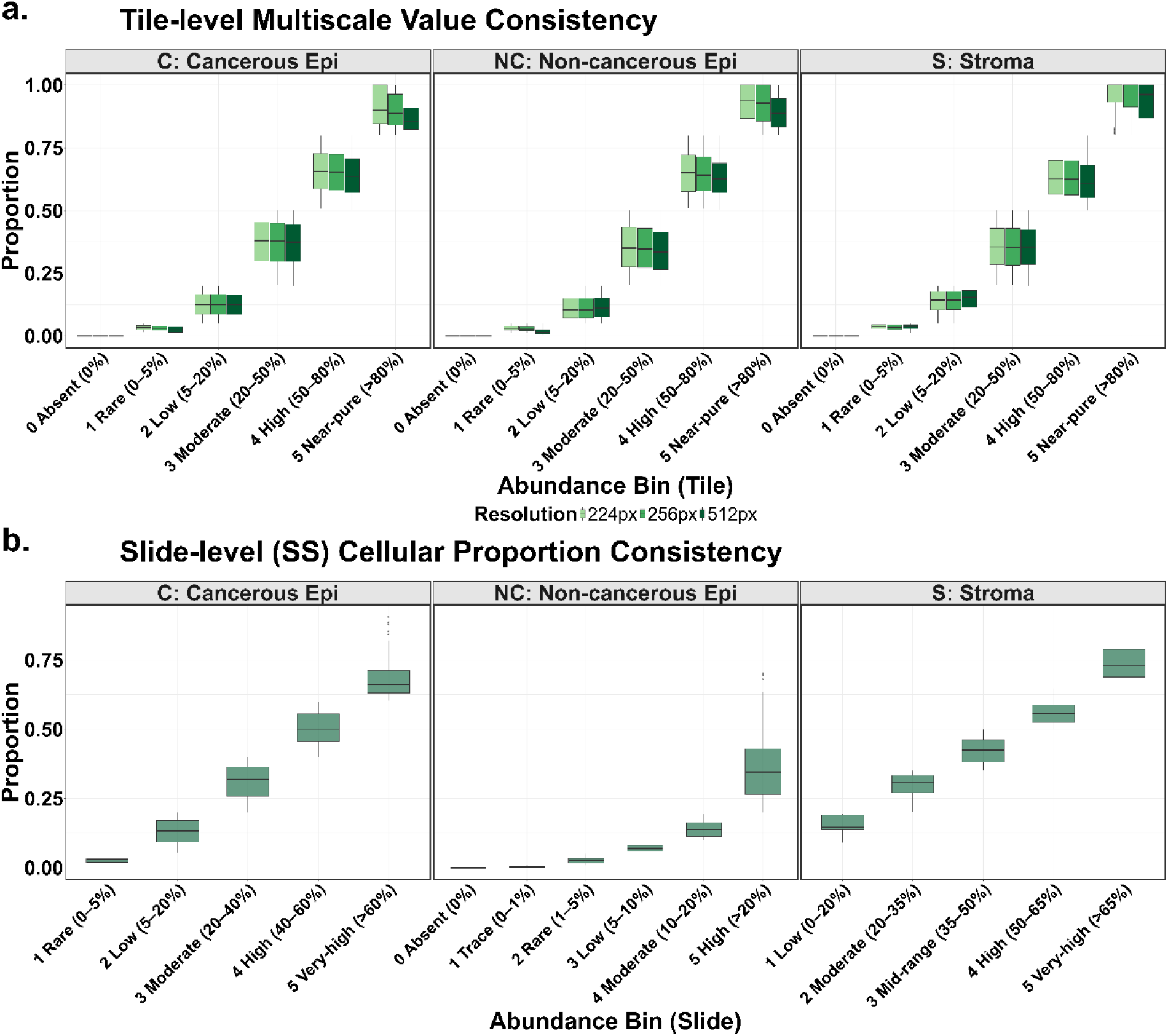
Multiscale Consistency of Cellular Composition across Image Resolutions. (a) Tile-level multiscale value consistency. Boxplots illustrate the proportional stability of three primary cellular modalities—Cancerous Epithelium (C), Non-cancerous Epithelium (NC), and Stroma (S)—across three distinct image resolutions: 224px, 256px, and 512px. Data are categorized into six abundance bins based on cellular proportions, ranging from ‘Absent (0%)’ to ‘Near-pure (>80%)’. The consistent median values across resolutions demonstrate the robustness of the multimodal labeling system against varying image scales. (b) Slide-level (SS) cellular proportion consistency across abundance bins. Distribution of whole-slide (SS) cellular proportions for each cell type, grouped by clinical abundance bins. Each dot represents a unique tissue slide (SS), confirming quantitative stability at the macro-level (slide) in addition to the micro-level (tile).

**Figure 5.**
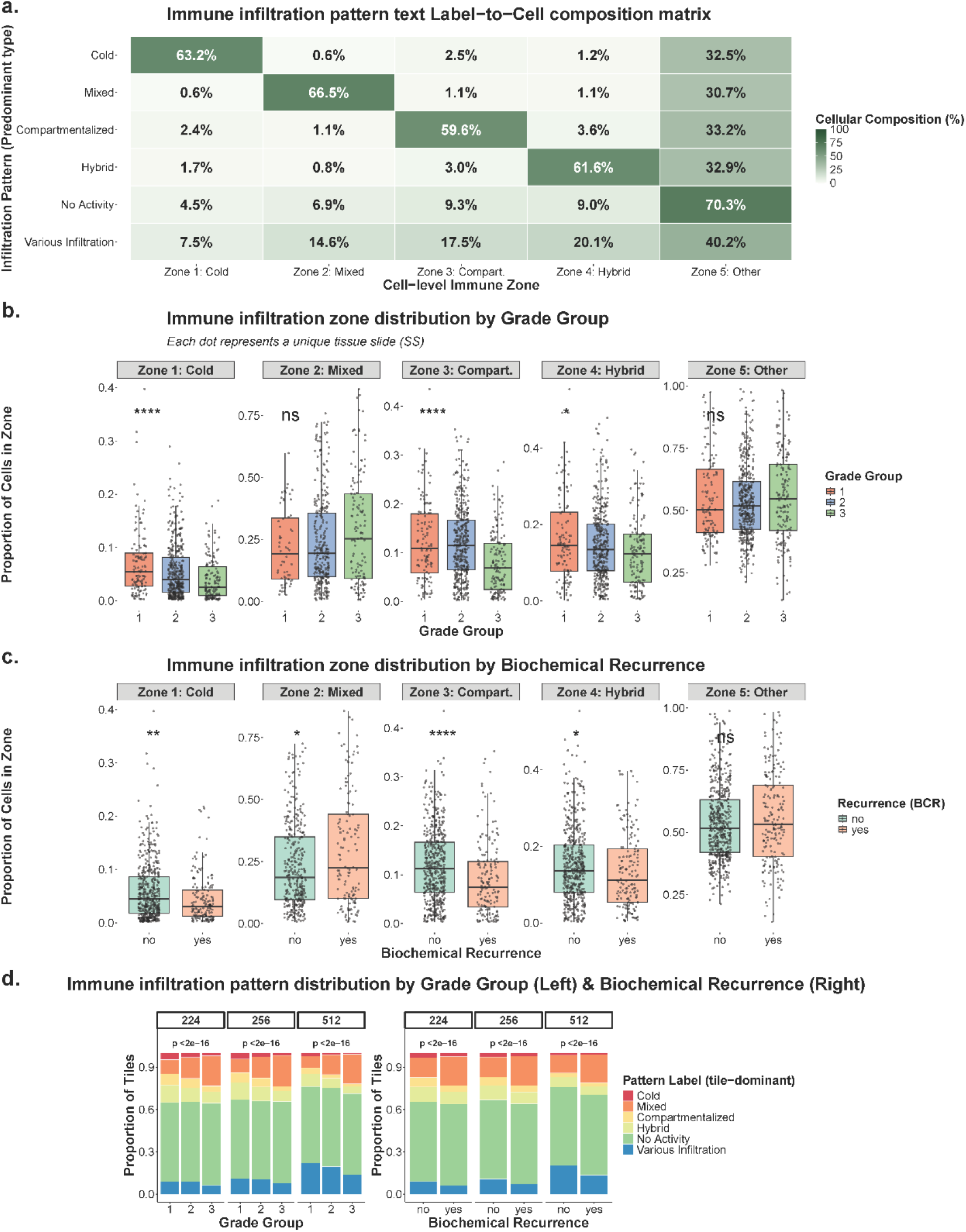
Technical Fidelity and Clinical Relevance of Multimodal Immune Infiltration Patterns. (a) Infiltration pattern text Label-to-Cell Composition Matrix. Heatmap illustrating the quantitative alignment between textual immune infiltration labels (e.g., Cold, Mixed, Compartmentalized) and the underlying cellular immune zone compositions (Zones 1–5). Percentages represent the mean cellular proportion of each zone within a given textual pattern, validating the descriptive accuracy of the multimodal labels. (b-c) Clinical validation of immune zones. Boxplots show the distribution of cellular proportions for each immune zone within each slide across (b) Grade Groups and (c) Biochemical Recurrence (BCR) status. Statistical significance was determined by Kruskal-Wallis (Grade) and Wilcoxon rank-sum (BCR) tests (****p < 0.0001, **p < 0.01, *p < 0.05, ns: not significant). (d) Distribution of tile-level predominant immune infiltration patterns across Grade Group and BCR. Stacked bar charts display the relative frequency of immune infiltration patterns across clinical categories. P-values derived from Chi-squared tests indicate a significant correlation between patterns and clinical disease progression.

**Figure 6.**
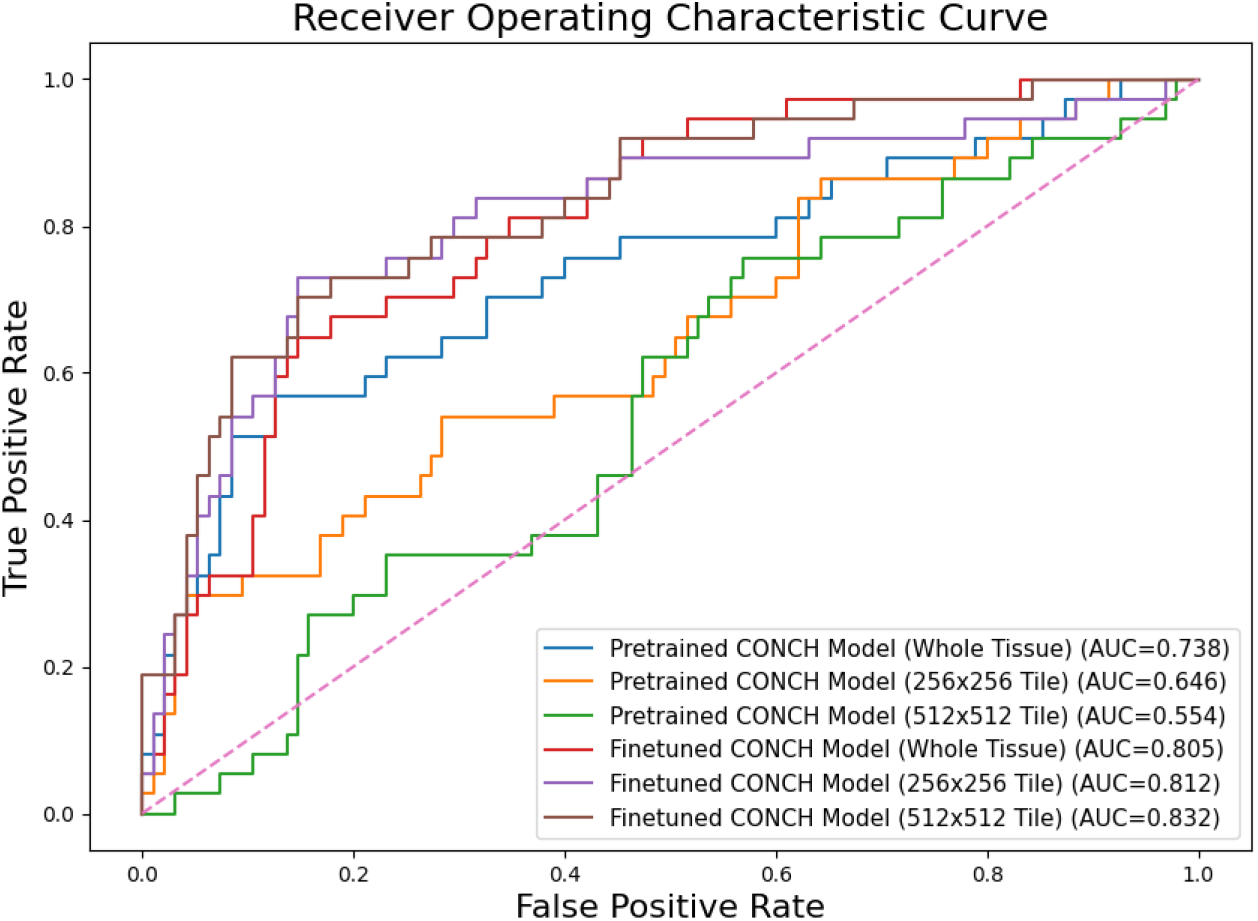
Comparative performance analysis of CONCH-based models using Receiver Operating Characteristic (ROC) curves. The consolidated ROC plot summarizes six model configurations spanning pretrained and fine-tuned CONCH models at both whole-tissue and tile resolutions. The pretrained CONCH whole-tissue model establishes the baseline, achieving an AUC of 0.738 using frozen foundation-model features. Pretrained CONCH tile-level models exhibit reduced discriminative performance, with AUCs of 0.646 at 256 px and 0.554 at 512 px, indicating limitations in out-of-domain representations. Fine-tuned models demonstrate substantial gains across all input scales: the fine-tuned whole-tissue model improves to an AUC of 0.805, while the fine-tuned 256 px and 512 px tile-level models further increase performance to 0.812 and 0.832, respectively. The overall progression from 0.738 to 0.832 AUC highlights the effectiveness of multi-scale fine-tuning, emphasizing both the value of domain-specific adaptation and the importance of larger spatial context for accurate prostate cancer prognosis.

### Single-cell Data Generation and Quality Control

Cellular segmentation^18^ was executed using a customized Fiji (ImageJ) plugin, utilizing DAPI and pan-cytokeratin signals to define epithelial and stromal boundaries along with individual cell ID, coordinates, area (pixel number), and expressions of markers. To ensure the technical integrity of the resulting dataset, we applied a QC pipeline at the cell level:

1. Nuclei Integrity: Epithelial cells were required to contain exactly one or two segmented nuclei.
2. Size Constraints: Sub-compartment areas (nucleus, membrane, cytoplasm) were restricted to 10–1,500 pixels. Total cell area thresholds were set at 50–3,500 pixels for epithelial cells and 30–1,500 pixels for stromal cells.
3. Registration Stability: We calculated a cyclic imaging stability score based on DAPI signal correlation across staining rounds. Only cells achieving a stability score ≥ 0.85 were retained.
4. Signal Normalization^19^: Batch effect correction and log2 transformation were applied to standardize intensity distributions across different imaging rounds.

After applying these criteria, a high-fidelity single-cell dataset of 724,742 epithelial cells and 552,702 stromal cells was established for analysis.

### Cell-type Classification using TOPAZ and CAT Models

The classification of individual cells was performed using a dual-model approach to capture both structural and phenotypic features. TOPAZ was primarily employed for the identification and classification of epithelial regions; it was specifically optimized to distinguish between Cancerous Epithelium (C) and Non-cancerous Epithelium (NC) based on their distinct glandular morphologies using p63 and CK5 markers.

In parallel, the CAT model was utilized for the classification of various immune cell populations. By the cell type definition of T cells and macrophages, T-helper cells (T_H_: CD3^+^CD4^+^), T-regulatory cells (T_reg_: CD3^+^CD4^+^FOXP3^+^), and macrophages (CD68^+^) were labeled. These designations were used to quantify immune-cell proximities to cancerous glands in downstream analyses. Due to the relatively low frequency of individual immune subtypes, these were aggregated into a single ‘Immune Cell’ category for subsequent spatial analysis and text caption generation to ensure statistical robustness. This comprehensive classification pipeline resulted in four primary cell categories: C, NC, Stroma (S), and Immune cells.

### Tissue (Cell-type) Map Generation

To visualize the spatial architecture of the tumor microenvironment, we generated pixel-level tissue (cell-type) maps by integrating the classification results with the spatial coordinates and area size of each cell from the single cell data. For each vH&E image, the identified cell phenotypes were mapped onto a spatial grid with a calculated area size, creating a multi-channel representation of the tissue. Blue/Orange/Green/Black dots represent C/NC/S/immune cells respectively. As immune cells are mostly from the S cells, they were encoded into triangle shape rather than a circle to distinguish from the S cells. The examples are shown in Figure 3b

### Multiscale Image Pre-processing

To support varied deep learning architectures and spatial reasoning tasks, whole-slide images (2040×2040 pixels at 20x magnification) were portioned into tiles across three spatial resolutions: 224×224, 256×256, and 512×512 pixels. This multiscale tiling approach maintains a constant pixel resolution while capturing different extents of spatial context—ranging from localized cellular distributions to broader glandular architectures within the tumor microenvironment (TME). Each data record in the repository is structured to maintain these varying scales and modalities, providing a robust foundation for multimodal model training.

### Text Caption Generation

Unlike datasets that rely on unstructured clinical reports or generative AI, our textual captions were generated through a rule-based mapping process grounded in quantitative single-cell data. This framework ensures 100% fidelity between the visual cellular architecture and the resulting descriptors, effectively eliminating linguistic hallucinations. The generation process involved four key components:

1. Cell count: Within each spatial scale (224px, 256px, 512px, and 2040px), the total number of cells was calculated directly from the synchronized single-cell data.
2. Cellular Abundance Mapping: Raw cell counts for each phenotype (C, NC, S) were converted into proportion levels within their respective scales. These proportions were then mapped to predefined categorical bins (e.g., ‘Absent’ (0%), ‘Rare’ (0-5%), ‘Low’ (5-20%), ‘Moderate’ (20-50%), ‘High’ (50-80%), ‘Near-pure’ (>80%)). The slide-level (SS) mapping utilizes specialized thresholds to reflect the broader tissue architecture as shown below in Table 2:
3. Dominant Immune Infiltration zone assignment: The distribution of individual cell types serves as the basis for calculating Immune Infiltration Zones. These patterns are derived from Euclidean distances between cancerous epithelium (glands) and immune cells, using a threshold of 15 µm. Every cell is assigned to one of five specific zones based on its proximity to the tumor:
  - Cold Zone: No immune cells are present within the 15 *µ*m threshold.
  - Mixed Zone: Over 5 immune cells are located within the cancerous glands.
  - Compartmentalized Zone: Immune cells are clustered at the periphery (edges) of cancerous glands within the threshold.
  - Hybrid Zone: Contains fewer than 6 immune cells within the glands while also exhibiting compartmentalized characteristics.
  - No Activity Zone: Cells that do not meet any of the criteria above indicate a lack of active immune infiltration.
  Dominant infiltration patterns are determined by calculating the proportion of these zones within the given scale. A pattern is labeled “Predominant” if it exceeds a specific proportion. For tile scales, the threshold is consistently ≥ 50%. However, for the slide-level (SS), the threshold for “No Activity” is increased to ≥ 80% to ensure precision at a macro scale, while other patterns retain a ≥ 50% threshold. If no single pattern meets these criteria, meaning that no pattern ≥ 50%, the slide is labeled as having “Balanced Infiltration” and “Various Infiltration” for tile scales.
4. Immune Infiltration Description Mapping: To provide natural language reasoning, the predominant patterns are mapped to specific biological descriptions, categorized by the spatial scale: For Whole Tissue (Slide-level, SS):
  - Predominantly No Activity: ≥ 80% of cells lack active immune infiltration.
  - Predominantly Cold: Minimal or no immune presence near glands across this slide.
  - Predominantly Mixed: Immune cells primarily found within glands on this slide.
  - Predominantly Compartmentalized: Immune cells mostly stay around gland edges for this slide.
  - Predominantly Hybrid: Both within-gland and around-gland infiltration dominate the slide.
  - Balanced Infiltration: No single pattern exceeds 50%; infiltration is more evenly distributed.
  For Tile Scales:
  - Predominantly No Activity: Most cells in this tile do not fall into an active immune zone category.
  - Predominantly Cold: Most glands show minimal or no immune presence (cold).
  - Predominantly Mixed: Immune cells primarily within glands (mixed).
  - Predominantly Compartmentalized: Immune cells cluster around gland edges (compartmentalized).
  - Predominantly Hybrid: Immune cells both inside and around glands (hybrid).
  - Various Infiltration: No single pattern exceeds 50%; multiple immune infiltration types coexist.

**Table 2.** Abundance bin rule for slide level.

| CANCEROUS EPI (C) | STROMA (S) | NON-CANCEROUS EPI (NC) |
| --- | --- | --- |
| ABSENT 0% | Absent 0% | Absent 0% |
| RARE (0-5%) | Low (0-20%) | Trace (0-1%) |
| LOW (5-20%) | Moderate (20-35%) | Rare (1-5%) |
| MODERATE (20-40%) | Mid-range (35-50%) | Low (5-10%) |
| HIGH (40-60%) | High (50-65%) | Moderate (10-20%) |
| VERY-HIGH (>60%) | Very-high (>65%) | High (>20%) |

**Table 3.** Comparative performance of pretrained and fine-tuned CONCH models across image-level and tile-level inputs, evaluated using Sensitivity, Specificity, Precision, Accuracy, F1-Score, and AUC.

| Model.Configuration | Sensitivity | Specificity | Precision | Accuracy | F1.Score | AUC |
| --- | --- | --- | --- | --- | --- | --- |
| Image-level Pretrained | 0.297 | 0.947 | 0.688 | 0.765 | 0.420 | 0.738 |
| Tile-level (256x256) Pretrained | 0.270 | 0.958 | 0.714 | 0.765 | 0.392 | 0.646 |
| Tile-level (512x512) Pretrained | 0.270 | 0.821 | 0.370 | 0.667 | 0.313 | 0.554 |
| Image-level Finetuned | 0.622 | 0.863 | 0.639 | 0.795 | 0.630 | 0.805 |
| Tile-level (256x256) Finetuned | 0.730 | 0.842 | 0.643 | 0.811 | 0.684 | 0.812 |
| Tile-level (512x512) Finetuned | 0.541 | 0.926 | 0.741 | 0.818 | 0.625 | 0.832 |

Finally, the structured text assembly phase synthesizes cell counts, categorical abundance bins, and predominant infiltration patterns and description into a unified, multiscale format. By using a structured template with these quantitative variables and their corresponding biological descriptions, we generate final captions through a purely deterministic process. This approach ensures that every component of the caption is directly anchored to a verifiable spatial-biological metric, as demonstrated by the representative examples in Figure 3c.

### Clinical Data Integration

To maximize the dataset’s utility for prognostic modeling, each slide was linked to clinical outcomes. This metadata includes Gleason Grade Groups and BCR status as well as the patient, slide information as shown in Figure 3d. The alignment between morphological patterns and clinical progression was rigorously maintained throughout the data processing pipeline to support the technical and clinical validations presented in Figure 5 and Figure 6.

## Technical Validation

Technical validation focuses on dataset consistency, label integrity, and practical usability for downstream analyses. The experiments below are provided as illustrative utility checks and reference baselines, rather than as the introduction of new modeling approaches.

### Technical Validation of Cell Type Classification and Immune Phenotyping

The fidelity of the Prostate-TriMod is anchored in the use of established and rigorously validated computational frameworks for cell classification and phenotyping.

The epithelial classification, which distinguishes cancerous (C) from non-cancerous (NC) cells, was performed using the TOPAZ model. The reliability of this model was previously established in Jung et al. (2025)^15^ using the same cohort through a concordance analysis with p63 expression, a definitive basal cell marker. The model achieves a high specificity of approximately 99% and a sensitivity range of 81.05%–81.89% compared to expert annotations, ensuring that the resulting spatial tissue maps and tri-modal captions are grounded in verifiable pathological features.

Similarly, immune cell phenotyping was conducted using the Cell Auto Training (CAT) model, a previously validated framework specifically designed for high-throughput phenotyping on the Cell DIVE™ platform. The CAT model provides a robust mechanism for characterizing individual cells based on multiplexed marker signals (CD3, CD4, CD8, CD68, and FOXP3). While we applied the CAT model to our cohort as a validated tool, its underlying framework has demonstrated high performance in literature, typically achieving overall accuracy exceeding 0.95 for major immune cell lineages. By leveraging these two high-performance models—one directly validated on this cohort and the other an established platform standard—we ensure that the immune-aware components of our dataset provide an accurate and biologically meaningful representation of the tumor microenvironment (TME).

### Multiscale Consistency and Structural Integrity

Since the dataset provides synchronized modalities across four spatial scales, we assessed whether cellular composition measurements remain consistent across varying spatial extents. A cross-scale analysis was performed focusing on the proportional distributions of three primary cell types: Cancerous Epithelium (C), Non-cancerous Epithelium (NC), and Stroma (S).

As shown in Figure 4a, tile-level distributions were compared across the 224px, 256px, and 512px resolutions using a uniform binning schema (from ‘Absent’ to ‘Near-pure’). The boxplots illustrate that the median cellular proportions are similar across these three scales, with concordance exceeding 95% between adjacent resolutions.

In Figure 4b, slide-level (SS) distributions at the 2040px scale are presented. To account for the broader tissue architecture represented at this scale, abundance bins with adjusted thresholds were defined for each cell type (e.g. ‘Very-high’ >60% for Cancerous Epithelium and >65% for Stroma). These calibrated bins provide a consistent framework for describing whole-tissue cellular composition across spatial scales.

### Biological Fidelity and Clinical Correspondence

The biological accuracy of our spatial phenotype maps and the corresponding rule-based captions were validated by analyzing their alignment with established clinical indicators, specifically Grade Groups and Biochemical Recurrence (BCR) status.

Figure 5a presents the Label-to-Cell Composition Matrix, which summarizes the relationship between the rule-based textual immune infiltration labels (e.g. ‘Cold’, ‘Mixed’) and the underlying immune infiltration zone distributions. For each tile, the mean proportions of cells assigned to the five immune infiltration zones were computed and grouped according to their corresponding textual labels. This matrix allows visualization of how textual immune infiltration categories relate to the cellular immune zone composition.

Figure 5b and 5c present the cellular composition of immune infiltration zones within each slide across Grade Groups and biochemical recurrence categories. For each slide, the proportions of cells assigned to the five immune infiltration zones were calculated and compared between clinical groups. Statistical tests (Kruskal-Wallis^20^ and Wilcoxon^21^) confirm that there are significant differences in zone composition across both Grade Groups and BCR status.

Finally, Figure 5d shows the distribution of tile-level predominant immune infiltration patterns across Grade Groups and BCR status. Each tile was assigned to a dominant immune infiltration label, and the proportions of these labels were compared between clinical categories. Chi-squared tests^22^ (*p* < 2*e* − 18) indicate a statistically significant association between tile-level immune infiltration labels and both Grade Group and recurrence status.

### Benchmarking Multimodal Utility for Prognostic Modeling

As shown in Figure 1c, to provide a reference benchmark for downstream prognostic modeling using Prostate-TriMod, we conducted a systematic benchmarking study using CONCH^23^, a multimodal foundation model pretrained on large-scale histopathology image–text pairs. CONCH is a transformer-based image encoder pretrained on histopathology image–text pairs using a contrastive objective. Here, CONCH is used solely as a feature extractor, with pretrained or fine-tuned image embeddings serving as inputs to downstream prognostic models for BCR prediction.

We focus on BCR prediction as a clinically relevant endpoint and summarize AUROC under standardized splits across multiple spatial scales and learning paradigms. Specifically, we evaluate frozen versus fine-tuned CONCH representations under two complementary strategies: whole-tissue–level classification using a multilayer perceptron (MLP), and tile-level modeling using a multiple instance learning (MIL^24^) framework. This design provides a controlled comparison of how spatial resolution, feature adaptation, and aggregation strategy relate to BCR prediction under a consistent dataset and evaluation protocol, and is intended as an illustrative baseline for reuse and benchmarking rather than a definitive performance claim. All evaluations use patient-level splits to avoid information leakage across tiles and to support a consistent benchmark protocol.

In the whole-tissue setting, each prostate tissue is treated as a single input that preserves global tissue architecture. Virtual H&E images are processed by the CONCH image encoder to extract a global tissue feature vector. These CONCH-derived features are then fed into an MLP classifier, which performs whole-tissue level BCR prediction. Fine-tuning allows the CONCH encoder to adapt to prostate-specific morphology, while the MLP learns a task-specific decision boundary in the learned feature space. As shown in Figure 6, fine-tuning yields higher AUROC than frozen features in this setting.

To capture fine-grained spatial heterogeneity that may be obscured at whole-tissue resolution, we further evaluate tile-level prognostic modeling using a MIL framework. Each tissue specimen is partitioned into non-overlapping tiles at 256px and 512px resolutions. Each tile is independently encoded by the CONCH image encoder, producing a set of tile-level feature vectors. These CONCH-derived tile embeddings are aggregated by a MIL module, which operates on bags of instances and outputs a single tissue-level prediction for BCR. Because supervision is available only at the tissue level, the MIL framework provides a mechanism to identify, and weight localized regions that are most predictive under the chosen endpoint. Fine-tuning of the CONCH encoder in this setting can further increase the discriminative utility of tile-level features.

## Usage Notes

The Prostate-TriMod dataset is structured to facilitate seamless integration into computational pathology and multimodal AI workflows.

### Potential Applications

This resource is designed to support a broad range of research directions in computational pathology and precision oncology, including but not limited to:

- Multimodal Foundation Model Training: Researchers can leverage the synchronized triplets for self-supervised or contrastive representation learning. The hallucination-free captions provide a reproducible foundation for pre-training or instruction-tuning VLMs, enabling models to learn hierarchical reasoning across cellular, and tissue scales.
- Causal and Interpretable Modeling: By combining the structured immune-zone labels with clinical outcomes, researchers can explore causal attributions—for instance, investigating whether specific compartmentalized immune patterns directly drive clinical risk or co-vary with Grade groups.
- Benchmarking and Methodological Development: The dataset serves as a high-fidelity benchmark for developing risk-stratification tools. This includes evaluating weak-to-strong supervision paradigms, where researchers can distill biological structures from our rule-based captions into neural captioners that generalize to external H&E datasets lacking multiplexed imaging.
- Decision-Support Prototypes: The integration of morphology and outcomes allows for the development of prototypes for interpretable AI tools that provide both prognostic predictions and visual/textual rationales (e.g., immune-zone maps) to assist clinical decision-making.

### Technical Implementation for Multimodal Modeling

For researchers utilizing this dataset for deep learning, the image-text-cell map triplets are provided in a synchronized format. To ensure high-fidelity training, the rule-based captions are deterministic and should be processed as fixed semantic descriptors rather than probabilistic natural language. Tile-level data records in the provided CSV files are indexed to match their corresponding PNG patches, allowing for direct batch loading in deep learning frameworks.

### Software Compatibility and Data Visualization/Processing

The dataset is provided in standard formats (TIFF/PNG for images and CSV for metadata) to ensure compatibility with diverse software environments.

- Image Visualization: It is recommended to use specialized digital pathology platforms such as QuPath (version 0.4+)^25^ or ImageJ/Fiji^26,27^ for visualization of TIFF formats as default OS image viewers may not be correctly rendered.
- Programming Environments: Structured metadata files are optimized for Python-based processing. Core libraries such as Pandas^28^ for metadata handling and PyTorch^29^ and TensorFlow^30^ for deep learning are recommended for loading the multiscale triplets.
- Reproducibility: Scripts used for data tiling and caption generation are available in the associated Zenodo repository^31^ to assist researchers in replicating the data processing pipeline.

## Data Records

Prostate-TriMod is deposited in the Zenodo repository under DOI https://doi.org/10.5281/zenodo.18209419 and is currently under restricted access during journal peer review. The public release will be activated upon acceptance of this manuscript. The data is organized into three primary modalities across four spatial scales, ensuring a structured and accessible format for multimodal analysis.

### Spatial Organization and File Formats

The repository is partitioned into scale-specific folders and master record files. Data is provided at three local patch resolutions (224×224, 256×256, and 512×512 pixels) and a whole-tissue scale (2040 pixels). As summarized in Figure 7, the dataset follows a strict naming convention and metadata structure:

**Figure 7.**
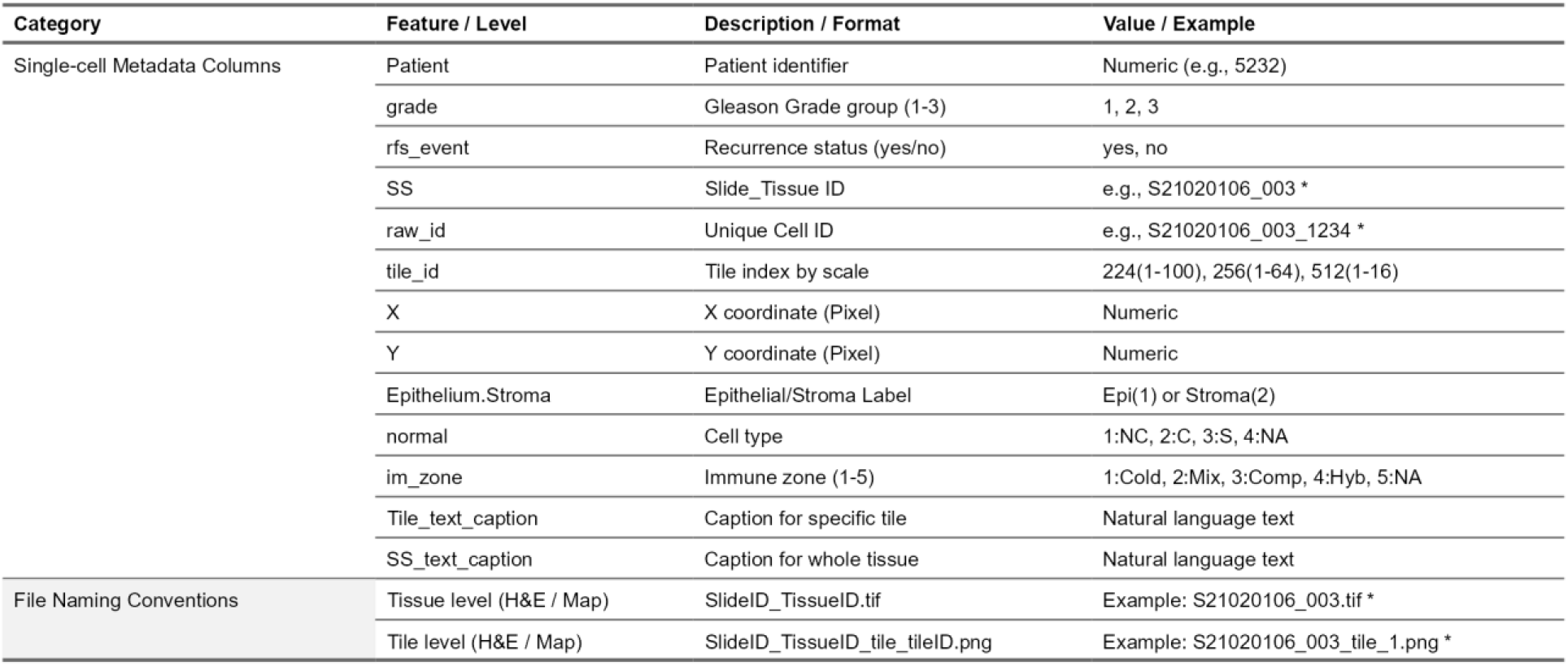
Detailed overview of metadata columns and phenotypic mapping logic. The table describes the standardized headers used across all CSV records, including phenotypic mapping and immune zone assignments. Abbreviations in the *im_zone* column are defined as follows: Mix: Mixed, Comp: Compartmentalized, Hyb: Hybrid. The asterisk (*) in the SS (Slide Tissue ID) example denotes identifiers assigned to whole-tissue records at the 2040 scale, distinguishing them from tile-level entries. * Note: TissueID must consist of exactly 3 digits (e.g., 001, 002, 003).

Within the repository, data is stored in the following formats:

- **Images (vH&E and tissue map)**: Multiscale patches are provided in PNG format (archived in ZIP files) and full-resolution whole-tissue images are provided in TIFF format.
- **Single-cell Metadata and Captions**: Our resource provides integrated CSV files (*single_cell_with_text_[scale]*.*csv*) for each resolution. These files contain individual cell IDs, x-y coordinates, assigned phenotypes (NC-Epi, C-Epi, Stroma, and Immune zones), and the corresponding rule-based text captions. The *Epithelium*.*Stroma* column provides a broad binary classification, distinguishing between epithelial (1) and stromal (2) compartments. The *normal* column identifies the primary structural phenotype for each cell: **Non-cancerous Epithelium (1: NC), Cancerous Epithelium (2: C), Stroma (3: S)**, and **Not Assigned (4: NA)**. While immune cells are explicitly visualized in the tissue map images as black triangles to provide architectural context, they are not provided as a separate cell-type label in the normal column. To prioritize spatial context over individual cells, immune-cell identities were utilized for internal zone computation and are released as spatial labels relative to the glands (*im_zone*). Specifically, the immune microenvironment is represented through the *im_zone* attribute (1: Cold, 2: Mixed, 3: Compartmentalized, 4: Hybrid, 5: NA). This approach ensures that immune information is captured as a spatial relationship between cancerous and immune cells, which serves as the foundation for the rule-based text captions.
- **Whole-tissue Master Records**: The *single_cell_with_text_whole_tissue*.*csv* file provides a comprehensive single-cell catalog for all slides at the 2040 scale. This master record contains the same column information as the multi-scale tile records, except for tile-specific indices.

Figure 7 presents detailed description of the metadata column description and its corresponding data format.

### Clinical Records

The dataset is accompanied by comprehensive de-identified clinical information integrated within the master and scale-specific CSV files in accordance with IRB guidelines. This includes Grade Groups, biochemical recurrence (BCR) status. This structure allows researchers to directly correlate local morphological features and spatial immune patterns with patient outcomes without the need for additional data merging.

## Data Availability

The Prostate-TriMod dataset has been deposited in Zenodo under DOI https://doi.org/10.5281/zenodo.18209419. The repository is currently under restricted access during journal peer review and will be made publicly available upon acceptance of this manuscript. The repository comprises three synchronized modalities: (1) multiscale virtual H&E images, (2) tissue map images, and (3) rule-based text captions. These data are provided across multiple spatial scales, including local tiles (224px, 256px, and 512px resolutions) and whole-tissue records (2040px scale). Comprehensive clinical metadata, including Grade Groups and biochemical recurrence (BCR) status, are also included. The data are released under the Creative Commons Attribution 4.0 International (CC BY 4.0) license. This release corresponds to Prostate-TriMod v1.0. Any future updates will be versioned via Zenodo.

## Funding

Research reported in this publication was supported by the National Cancer Institute of the National Institutes of Health (NIH) under Award Number R01CA249899 (PM). Funder website: https://www.cancer.gov/. The funders had no role in study design, data collection and analysis, decision to publish, or preparation of the manuscript. The content is solely the responsibility of the authors and does not necessarily represent the official views of the National Institutes of Health.

## Author Contribution

KJJ: Conceptualization, Validation, Visualization, Writing – Original Draft Preparation. JQ: Benchmarking, Writing – Review & Editing. SC: Data curation, Investigation, Writing – Review & Editing. EM: Investigation, Validation, Writing – Review & Editing. CC: Investigation, Validation, Writing – Review & Editing. SG: Writing – Review & Editing. RW: Writing – Review & Editing. JDB: Resources, Writing – Review & Editing. FG: Supervision, Writing – Review & Editing. RM: Supervision, Writing – Review & Editing. PM: Supervision, Funding Acquisition, Writing – Review & Editing.

## Code Availability

The source codes (R, ipynb) for the dataset construction pipeline—specifically the multiscale patch extraction (tiling), and rule-based captioning logic—is available in the ‘Script’ folder of the Zenodo repository (https://doi.org/10.5281/zenodo.18209419) and will be made publicly available upon acceptance of this manuscript. While the specific implementations of the TOPAZ and CAT pre-processing models are not included to maintain data security, the resulting high-fidelity phenotypic metadata (cell IDs, coordinates, and assigned zones) is fully provided in the shared CSV files to ensure complete reproducibility of the dataset’s structure. Researchers seeking to apply these specific pre-processing models to new cohorts should refer to the methodologies validated in Jung et al. (2025)^15^ and Santamaria-Pang et al. (2019)^17^.

## Notes

### Competing Interest Statement

The authors have declared no competing interest.

